# Machine Learning-Based Tumor Segmentation and Classification Using Dynamic Optical Contrast Imaging (DOCI) for Thyroid Cancer

**DOI:** 10.1101/2025.02.03.636242

**Authors:** Tyler Vasse, Yazeed Alhiyari, Lauran K. Evans, Ramesh Shori, Maie St. John, Tuan Vo-Dinh

**Affiliations:** Fitzpatrick Institute of Photonics, Duke University; Durham, NC 27708, United States of America; Department of Biomedical Engineering, Duke University; Durham, NC 27708, United States of America; Department of Head and Neck Surgery, David Geffen School of Medicine at University of California; Los Angeles, CA 90025, United States of America; Head and Neck Cancer Program at University of Los Angeles; Los Angeles, CA 90025, United States of America; Department of Chemistry, Duke University; Durham, NC 27708, United States of America; Jonsson Comprehensive Cancer Center at University of Los Angeles Medical Center; Los Angeles, CA 90025, United States of America

## Abstract

Thyroid cancer presents significant diagnostic challenges due to its complex anatomy and diverse tissue types. This study leverages Dynamic Optical Contrast Imaging (DOCI), a label-free, real-time imaging technology, with machine learning to enhance tumor detection and segmentation. Using 23 DOCI filters, we applied Principal Component Analysis (PCA) for dimensionality reduction, k-nearest neighbors (k-NN) for classification, and U-Net models for segmentation. The approach achieved high accuracy in distinguishing tissue types, with PCA enabling clear clustering, k-NN classifying normal, follicular, and papillary tissues, and U-Net models achieving 96.34% and 92.02% accuracy for papillary and follicular segmentation, respectively. Filter importance analysis reduced input dimensionality without significantly compromising performance, highlighting the potential for optimized imaging protocols. These findings demonstrate DOCI’s utility in improving diagnostic accuracy and tumor characterization in thyroid cancer and beyond, offering a foundation for personalized treatment planning and surgical precision.

## 1. Introduction

Thyroid cancer is a significant health concern, representing the most common endocrine cancer worldwide continues to increase each year [1, 2]. This malignancy arises from the thyroid gland, a butterfly-shaped organ in the anterior neck that plays a crucial role in regulating metabolism through thyroid hormone production [3]. Thyroid cancer encompasses several subtypes, including papillary, follicular, medullary, and anaplastic, each with distinct characteristics, treatments, and prognoses [3, 4]. Over recent decades, the incidence of thyroid cancer has been steadily increasing, largely attributed to advancements in diagnostic techniques and increased surveillance [5, 6]. This rise is particularly evident in papillary thyroid cancer, which has seen the most substantial growth in cases [6]. While the exact causes of thyroid cancer remain unclear, risk factors include exposure to ionizing radiation, family history, and certain genetic conditions [3, 7].

The complex anatomy of the neck region and the proximity of the thyroid gland to vital structures make accurate diagnosis and treatment challenging [8]. This complexity further complicates the differentiation between thyroid cancer and benign thyroid nodules, which affect up to 50% of individuals over the age of 50 [8].

Traditional diagnostic approaches for thyroid cancer often involve a combination of physical examination, blood tests, imaging studies, and fine-needle aspiration biopsy. The fine-needle aspiration (FNA) biopsy is considered the “gold standard” method for diagnosing thyroid nodules [9]. However, these methods can sometimes lead to inconclusive results or unnecessary surgeries for benign nodules [6]. In fact, 70-80% of thyroid nodules with indeterminate FNA results are ultimately found to be benign upon histological analysis of surgical specimens [10]. Additionally, FNA cannot provide spatial information about the specimens. Such spatial information is crucial during surgical removal, as positive tumor margins have significant prognostic implications [11]. Currently, tumor margins are assessed through Hematoxylin & Eosin (H&E) staining after surgical resection, which cannot be performed during surgery to provide real-time feedback.

As such, there is a pressing need for advanced imaging techniques that can provide more accurate, spatially-resolved, real-time and detailed information about thyroid nodules and potential malignancy and subtypes [8]. The development of novel imaging modalities has the potential to revolutionize the management of thyroid cancer by enabling earlier detection, more precise staging, and better-informed treatment decisions, and enable real-time tumor margin assessment [12].

Our team has previously developed and described Dynamic Optical Contrast Imaging (DOCI), an innovative imaging technology that captures time-dependent measurements of tissue autofluorescence [13-17]. In fresh ex vivo studies, we demonstrated that the DOCI system can effectively distinguish HNSCC from adjacent normal tissue and accurately localize tumors [13]. This imaging method operates in real time, does not require the use of contrast agents, and provides a broad field of view during surgical procedures. By leveraging the fluorescence lifetime of various endogenous fluorophores within the tissue, DOCI generates a distinct molecular map to aid in identifying cancer margins. These margins are then validated against histological slides, which serve as the reference standard. Earlier studies have explored methods to statistically analyze DOCI images for tissue classification, employing approaches that range from basic logistic regression to advanced artificial neural networks (ANNs) [18].

This paper presents an analysis method to detect cancerous regions from DOCI images in histopathologic slides, combining advanced data acquisition with cutting-edge machine learning integrated techniques to enhance diagnostic accuracy. Our approach analyzes multi-wavelength DOCI data collection using Principal Component Analysis (PCA) for dimensionality reduction, k-nearest neighbors (k-NN) for tissue classification, and a U-Net-based architecture for precise tumor segmentation [13, 19]. We identified the most informative DOCI wavelengths using feature selection, enhanced model robustness through extensive data augmentation, and comprehensively characterized the tumor via, multi-filter. By integrating classification and segmentation processes, our technique achieves high accuracy in distinguishing between normal and cancerous tissues, including follicular and papillary types [20]. This innovative approach not only improves diagnostic precision but also provides detailed insights into tumor characteristics, potentially transforming treatment planning and patient care in head and neck oncology.

## 2. Methods

### 2.1 Data Collection and Preparation

Healthy and cancerous thyroid tissue samples (n=73) were collected in the operating room and processed through the pathology department according to standard of care. These samples were obtained to investigate their optical properties and assess the ability of DOCI to distinguish between cancerous and non-cancerous tissue regions. After the pathology department processed the samples, the corresponding tissue blocks were then retrieved and 5 micron unstained slides were cut from those specimens to first DOCI image, then stain with H&E for comparison. The imaging utilizing DOCI occurred at 23 distinct wavelengths, ranging from DOCI 1 to DOCI 23 (Supplementary Table 1).

**TABLE 1.**
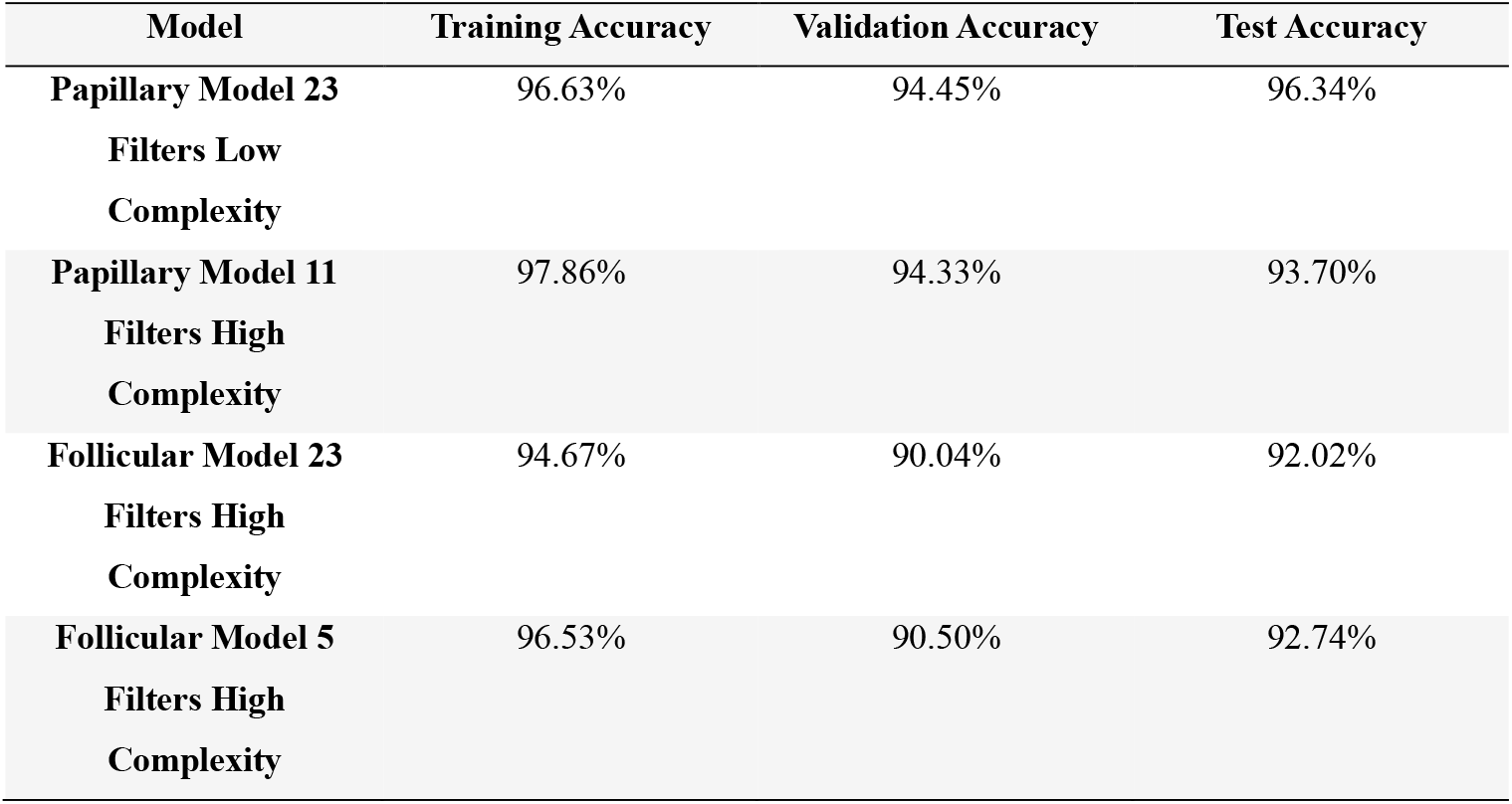
MODEL PERFORMANCE. Performance metrics for training, validation, and test sets for both models using all 23 filters and a subset of the most important filters. Accuracy was measured using Tensorflow’s built-in accuracy metric.

Each stained tissue sample was digitized and then annotated by a head & neck trained pathologist to identify whether the sample contained cancer or was entirely free of cancerous regions. Samples that had no cancer present were designated as “Normal.” For samples identified as cancerous, the pathologist further classified the cancer type into one of three categories: Follicular, Anaplastic, or Papillary. Medullary thyroid carcinoma samples were not included in this study due to lack of tissue samples.

To provide spatial localization of the cancerous regions, the pathologist created detailed annotations, which were provided in XML files. These annotations outlined the exact areas within the tissue samples that corresponded to cancerous regions, as well as non-cancerous regions. This allowed for precise mapping of DOCI data to tumor locations and enabled pixel-level separation of cancerous and non-cancerous regions. Normal samples provided a critical baseline for comparison, ensuring the model could differentiate between healthy tissue and various cancer types, while the inclusion of spatial annotations facilitated a deeper understanding of the optical variations within cancerous tissues. This comprehensive dataset formed the foundation for training and evaluating the classification and segmentation models.

## 3. Results

### 3.1 Preprocessing and Dimensionality Reduction

To ensure robust analysis of the high-dimensional DOCI data, Principal Component Analysis (PCA) was applied as a pre-processing step to reduce the dataset’s dimensionality before further classification, a strategy that has been previously utilized in similar contexts with limited data [21]. PCA was specifically chosen because it is an unsupervised learning technique, which minimized overfitting risks while effectively identifying patterns in the data [22].

For each pixel in the tissue image, the DOCI values across the 23 wavelengths were compiled into a 23-dimensional vector. Background pixels were identified and excluded using tissue-specific masks. The remaining tissue pixel vectors for each sample were used as input to PCA. This per-pixel PCA enabled the analysis to be independent of the varying sizes and shapes of extracted tissue samples, a challenge that conventional convolutional models often struggle to address. This transformation projected the 23-dimensional DOCI data into a two-dimensional space, enabling visualization and comparison of tissue types (Supplementary Fig. 1). After performing PCA, the principal component values of all valid pixels were averaged for each sample, producing a unique PCA profile for every tissue sample. This process ensured the analysis captured the overall DOCI patterns specific to each tissue type.

The results of the PCA analysis are visualized in Fig. 3A, which displays the distribution of tissue samples in the PCA space, color-coded by tissue type. Normal samples clustered distinctly from cancerous samples, with Follicular and Papillary samples forming separate groups. Anaplastic samples, known for their heterogeneous DOCI profiles, appeared interspersed among both cancerous clusters. This effective separation, particularly for Normal, Follicular, and Papillary samples, highlights the utility of PCA in distinguishing tissue types based on DOCI data. The amount of variance captured by each of the principal components was analyzed and graphed (Fig. 3B). The first two principal components, which together accounted for 79.9% of the variance, underscore the dominance of these two dimensions in capturing the most critical features of the DOCI data, enabling a highly efficient and interpretable reduction of the dataset’s complexity while preserving its most informative patterns. Furthermore, the clear clustering provided the basis for downstream classification tasks using k-nearest neighbors (k-NN), enabling accurate tissue type identification while accounting for the heterogeneity of Anaplastic samples.

**Fig. 1.**
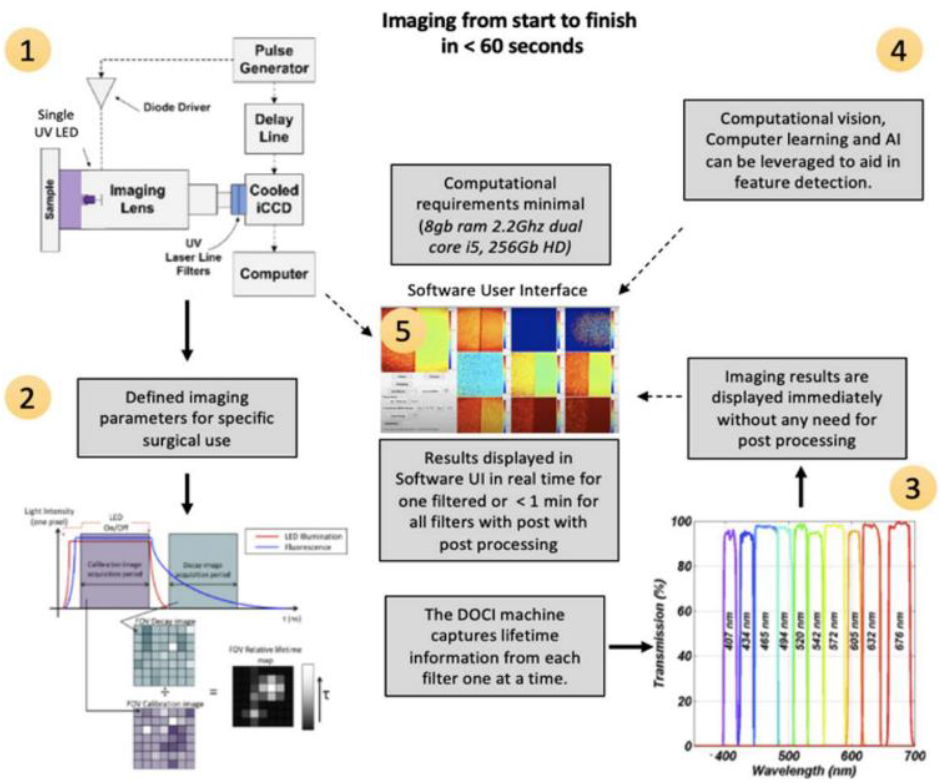
(1) Diagram illustrating the DOCI system architecture. (2) The image generation process begins with predefined imaging parameters tailored for specific surgical applications. (3) Images are acquired at the corresponding wavelengths. (4) AI-powered and computer-assisted feature detection can be integrated into the software interface. (5) The user-friendly software interface allows image or video acquisition in just three clicks, with immediate display of imaging results, eliminating the need for post-processing.

**Fig. 2.**
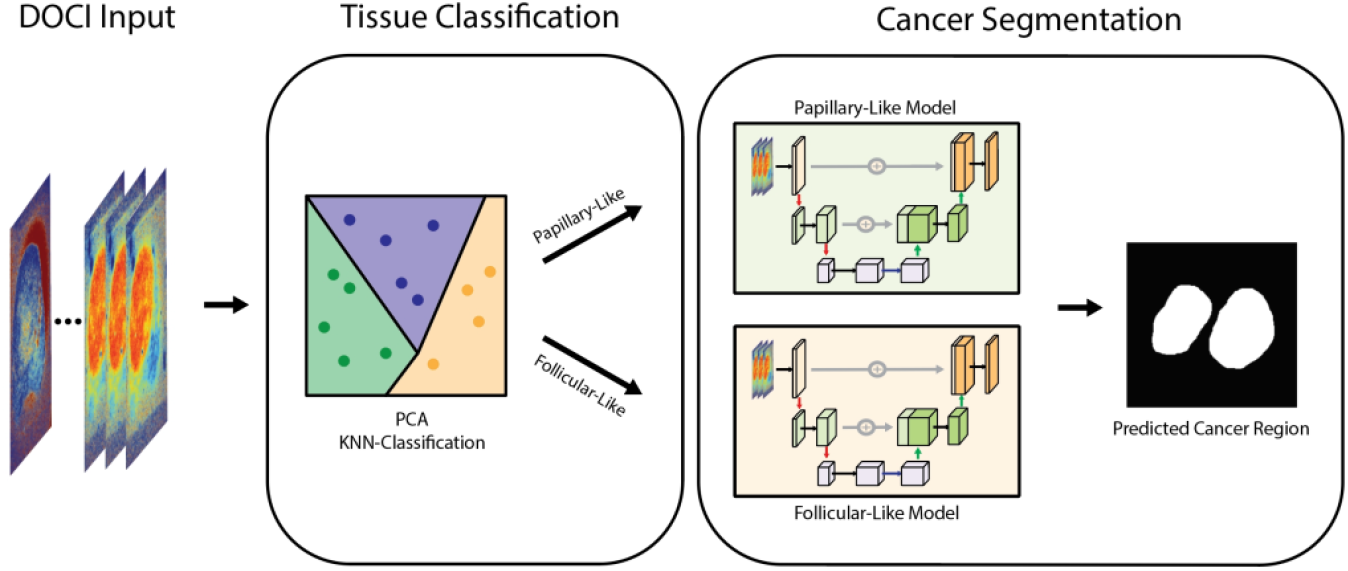
Overview of the DOCI-based tissue classification and cancer segmentation pipeline. The pipeline begins with Dynamic Optical Contrast Imaging (DOCI) input across 23 spectral filters. Principal Component Analysis (PCA) and k-nearest neighbors (k-NN) classification are used to classify tissue into Papillary-like or Follicular-like categories. These classifications guide segmentation models, where U-Net architectures tailored for each tissue type predict cancerous regions, producing precise masks for tumor localization.

**Fig. 3.**
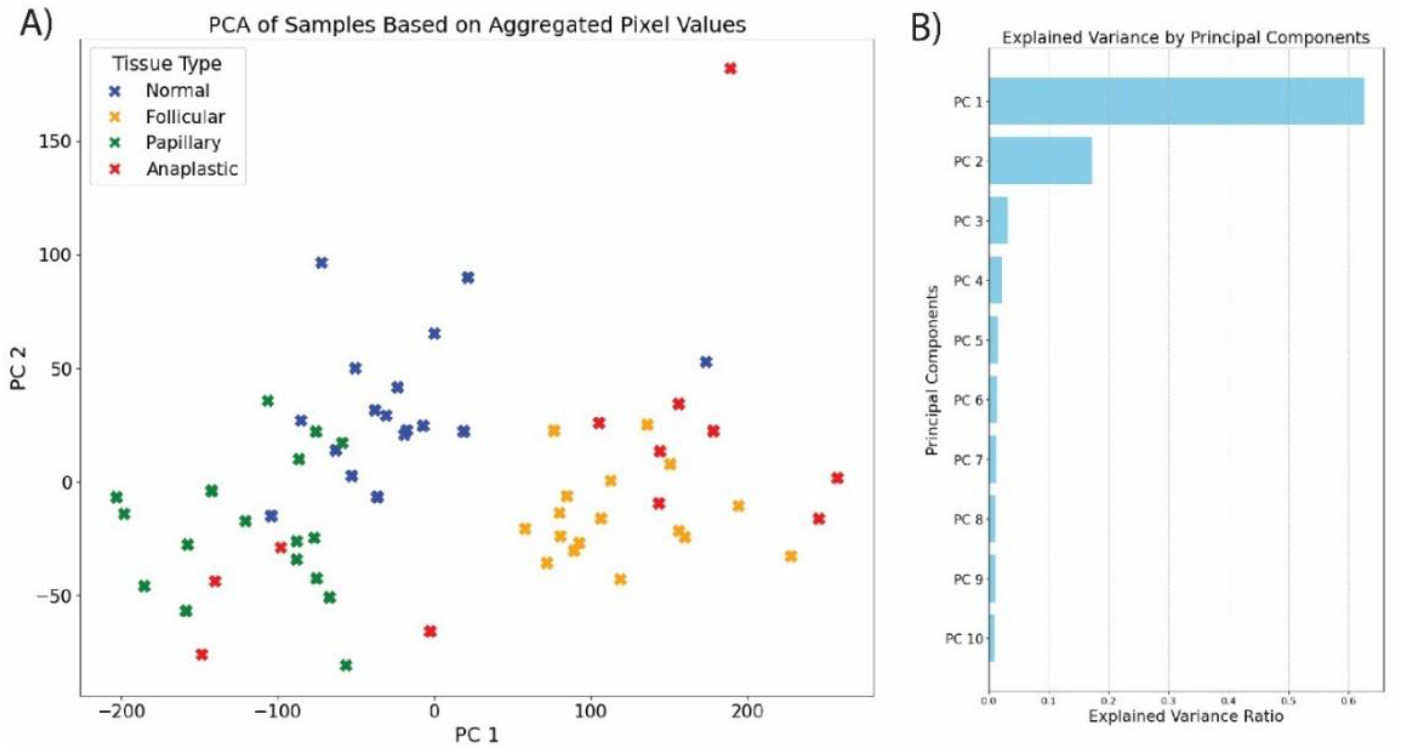
(A) PCA of tissue samples based on DOCI data aggregated from 23 wavelengths, displaying distinct clustering of Normal (blue), Follicular (orange), Papillary (green), and Anaplastic (red) tissue types. Normal, Follicular, and Papillary samples form clearly defined clusters, while Anaplastic samples exhibit heterogeneous distributions. (B) Bar plot illustrating the explained variance ratio of the principal components derived from PCA on the DOCI dataset. The first principal component (PC1) explains the majority of the variance (62.7%), followed by the second principal component (PC2) accounting for 17.2%, together capturing 79.9% of the total variance.

### 3.2 K-NN for Tissue Classification

To classify tissue samples into Normal, Follicular, and Papillary categories, a k-nearest neighbors (k-NN) algorithm was applied to the PCA-transformed data. Due to their overlapping distributions with the two other cancerous classes, Anaplastic samples were excluded from the primary classification step. Instead, their accuracy was assessed separately, where Anaplastic samples were considered correctly classified if they were categorized as anything other than Normal.

We employed 8-fold cross-validation to optimize the number of neighbors (*K*) for the k-NN (Fig. 4B). For each fold, the k-NN model was trained with *K* values ranging from 1 to 10. Combined accuracy was calculated by considering both the classification accuracy of non-Anaplastic samples in the validation set and the success rate of classifying Anaplastic samples as non-Normal. These metrics were averaged across all folds, with the standard deviation of accuracy recorded to assess variability. Additionally, test set accuracy was calculated for each *K* value, including Anaplastic samples, to evaluate the model’s generalization to unseen data. The cross-validation results revealed that training accuracy stabilized as *K* increased, while test accuracy remained consistently high across a range of values. The optimal *K* was determined to be K=7 as it was the highest average cross-validation accuracy during training (Fig. 4B)

**Fig. 4.**
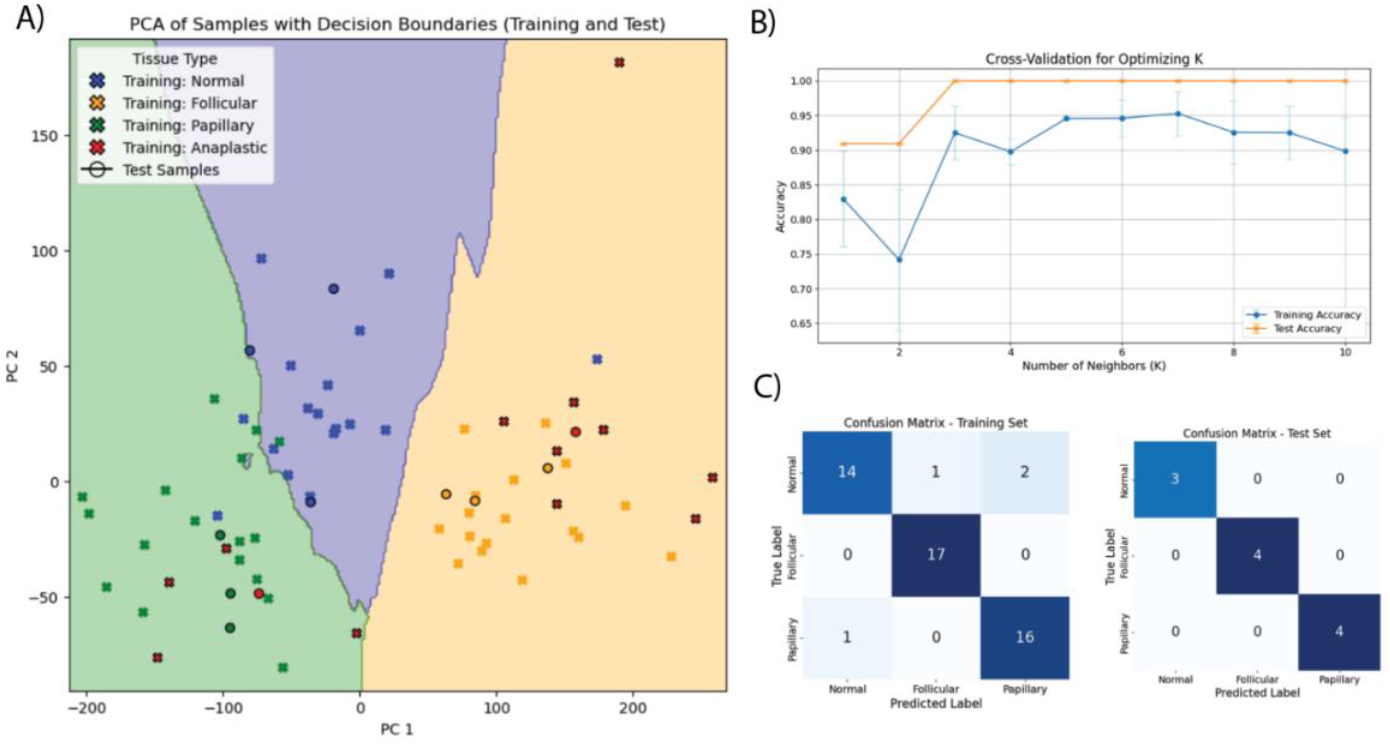
(A) Decision boundaries generated by the k-nearest neighbors (k-NN) classifier in PCA-transformed space, trained to separate Normal, Follicular, and Papillary tissue types. Training samples are marked with solid shapes, while test samples are shown as outlined markers. Anaplastic samples, highlighted separately, span multiple clusters due to their variability, further validating the heterogeneity of this tissue type. (B) Cross-validation results for optimizing the number of neighbors (K) in the k-NN classifier. Training accuracy (orange) and test accuracy (blue) are plotted as a function of K, with error bars representing standard deviations across folds. (C) Confusion matrices illustrating the classification performance of the k-NN model on the training set (left) and test set (right) for the tissue samples.

To illustrate the decision-making boundaries of the classifier, decision regions were visualized in PCA space using *K*=7 (Fig. 4A). This plot included both training and test samples, with Normal, Follicular, and Papillary samples represented by distinct markers and colors, and Anaplastic samples highlighted separately. The decision regions demonstrated clear separation between Normal, Follicular, and Papillary samples, while Anaplastic samples were dispersed across both cancerous class boundaries due to their heterogeneous nature. This visualization further emphasized the effectiveness of the PCA-based classification and the ability of the k-NN classifier to delineate tissue types. This combined approach, integrating PCA for dimensionality reduction, k-NN for classification, and cross-validation for hyperparameter optimization, effectively separated Normal, Follicular, and Papillary samples while also successfully identifying Anaplastic tissues as cancerous.

### 3.3 Data Augmentation and Model Architecture

Following the limited success of using joint classification and segmentation models with mask-based voxel approaches, the decision was made to build two separate U-Net architectures tailored for specific tissue classifications derived from PCA analysis. These models were designed to leverage the distinct clustering observed in the PCA-transformed space, grouping samples into Papillary-like and Follicular-like categories. Papillary-like includes all samples classified as Papillary in the k-NN, including Anaplastic samples, while Follicular-like includes those classified as Follicular, also including Anaplastic samples. Each U-Net was designed to optimize segmentation for its corresponding tissue type, addressing the unique challenges posed by the heterogeneity within each category. The models were trained using DOCI samples and their corresponding pathologist-identified annotations (masks), which served as ground truth.

To address the small dataset size, extensive data augmentation was performed on copies of the DOCI samples and their corresponding masks. Augmentation techniques included horizontal flipping, random zooming, resizing, and random noise addition. All images and masks were normalized before inputting into the model.

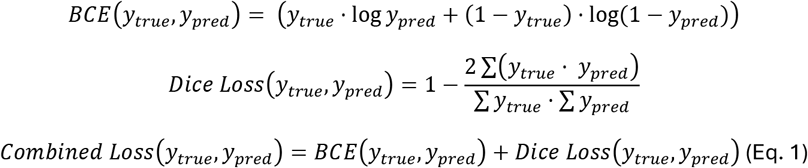

The architecture of the U-Net employed for this task consisted of encoder and decoder blocks to progressively extract features, reduce the spatial dimension, and subsequently reconstruct the spatial resolution (Fig. 5A). Each block incorporated convolutional, batch normalization, and ReLU activation layers. Instead of using max-pooling for down-sampling, 2×2 strided convolutions were implemented, ensuring smoother spatial reduction while retaining contextual information. The decoder utilized transposed convolutions for up-sampling, paired with skip connections to preserve fine-grained spatial features by directly linking corresponding encoder and decoder layers. The model incorporated a combined loss function that balanced Dice Loss and Binary Cross-Entropy Loss (BCE) (Eq. 1), ensuring the model performed well on both segmentation quality and pixel-level classification. The model was trained using a 55/30/15 Training/Validation/Test data split. Each tissue type was split individually to ensure all groups had a representative set of images for that type.

**Fig. 5.**
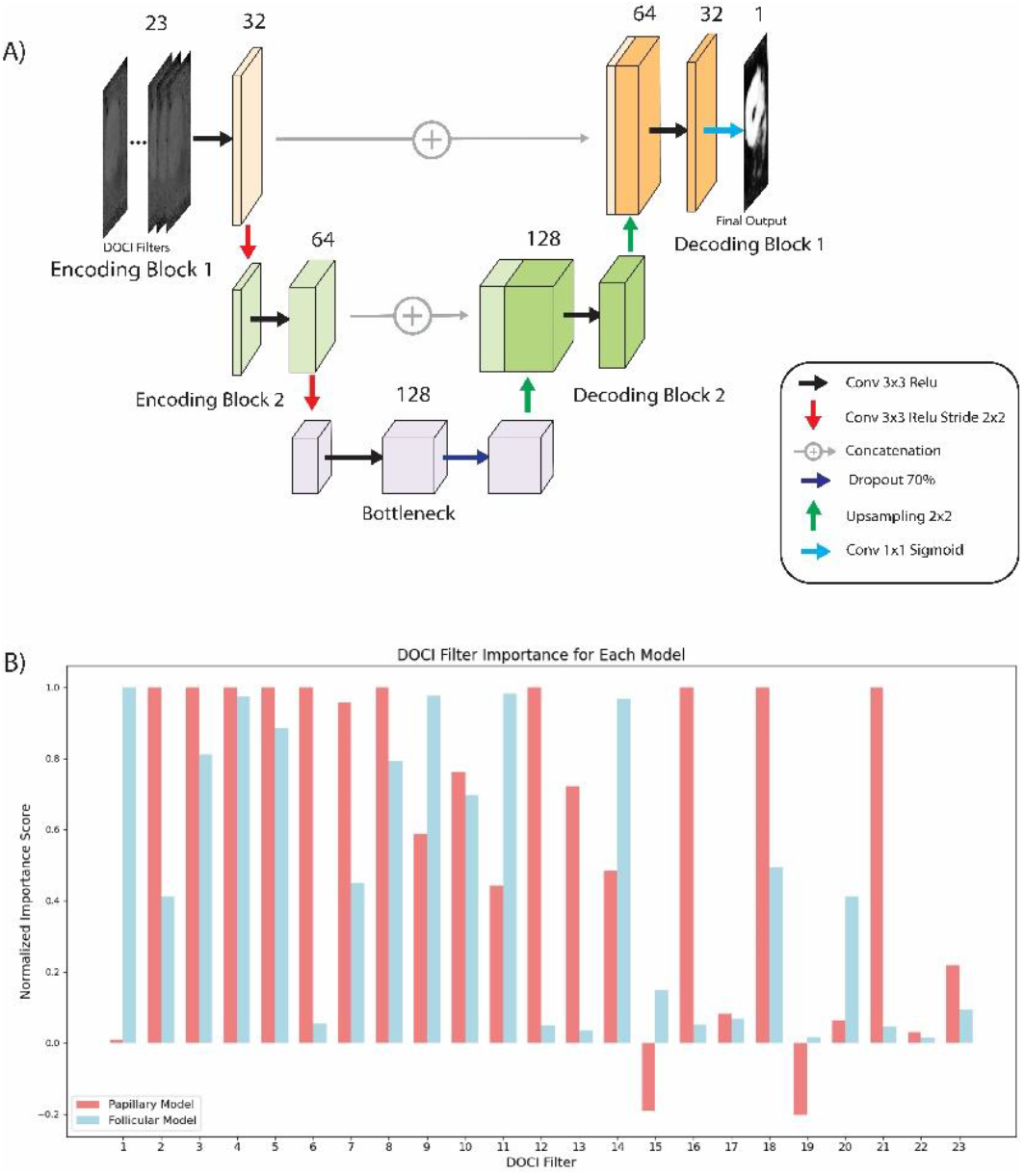
(A) Diagram of the U-Net architecture used for segmentation, including encoding and decoding blocks with skip connections, convolutional layers, and dropout. (B) Filter importance analysis for the Papillary (red) and Follicular (blue) models, showing the normalized impact of each DOCI filter on model performance.

**Fig. 6.**
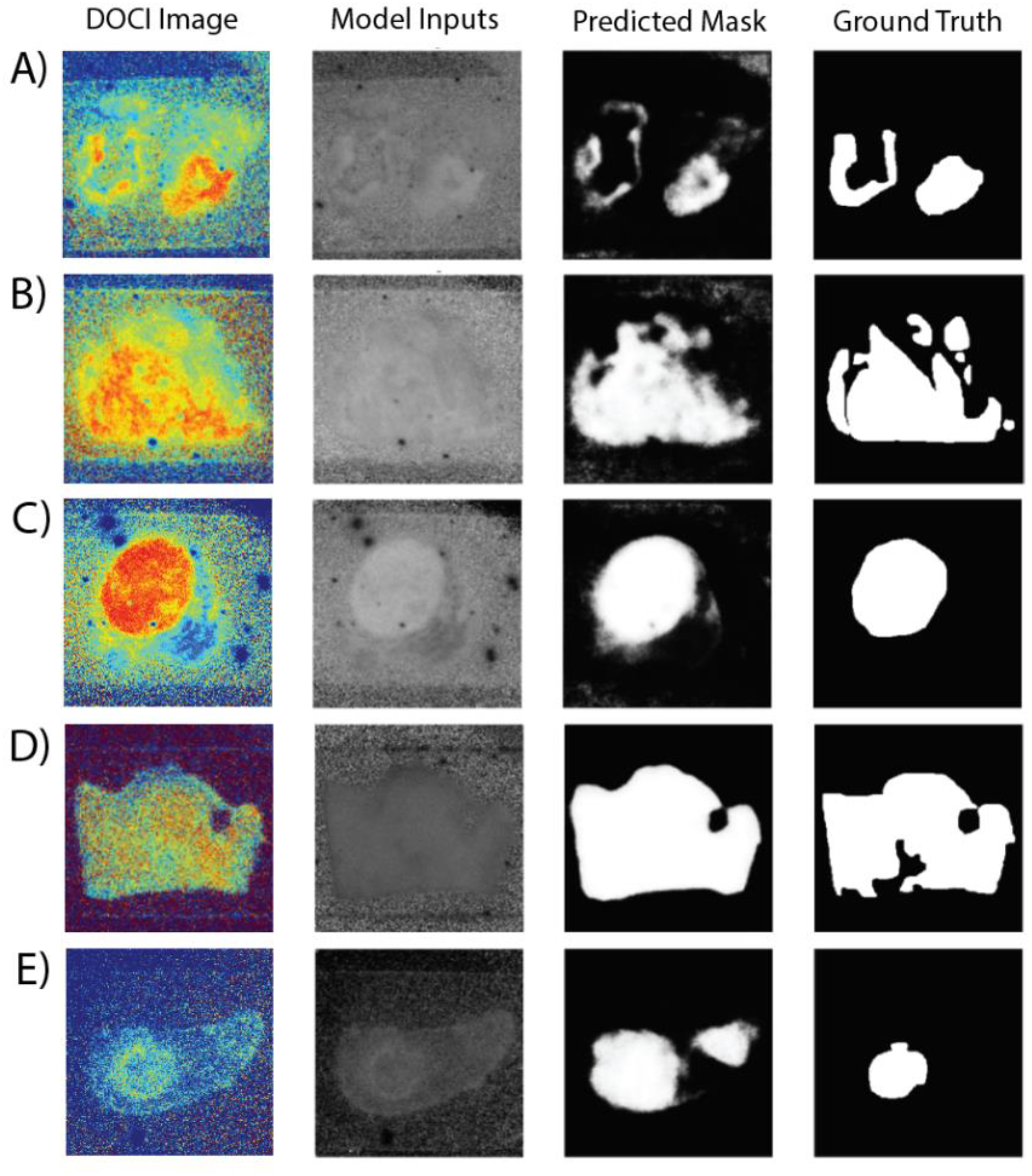
Test set results from the two segmentation models utilizing all 23 DOCI filters. Panels A-C display segmentation outcomes generated by the Papillary model, while panels D-E showcase results produced by the Follicular model. Images in the DOCI Image column have been converted to RGB for visualization purposes.

### 3.4 Training and Evaluation

In both the original Papillary and Follicular models, the full set of 23 DOCI filters was utilized as input, capturing the comprehensive spectral information from the tissue samples. These models were trained on a combination of images identified by the PCA analysis as Papillary or Follicular tissues, with Normal samples included as negative controls to enhance model robustness. Despite their shared input structure, the complexity of the tissue types influenced the design and performance of each model. Through an iterative optimization process, it was determined that the Papillary model required only two encoding and decoding blocks to effectively capture its more distinct and localized features. In contrast, the Follicular model, characterized by more abstract and dispersed features, required an additional encoding and decoding block to improve feature representation and segmentation accuracy.

The Papillary model achieved an impressive test accuracy of 96.34%, demonstrating its ability to reliably identify Papillary tissue regions. On the other hand, the Follicular model achieved a slightly lower accuracy of 92.02%, reflecting the increased complexity and variability within this tissue type. Despite these challenges, both models provided strong segmentation performance across their respective tasks.

To investigate the biological relevance of each filter, an analysis of filter importance was conducted for both models. This was achieved by systematically zeroing out individual filters from the validation set and passing the modified input through the model. The impact on performance, measured through changes in Intersection over Union (IoU), was recorded and visualized (Fig. 5B).

Using a cutoff of 0.90, the critical filters were identified for each model, and the models were retrained using only these top filters. Interestingly, the performance of the Papillary model declined significantly with this reduced input, highlighting its dependence on the full spectral information. Conversely, the Follicular model maintained its accuracy despite the reduced input, suggesting that the selected filters effectively captured the most relevant features for this tissue type (Table 1). This finding underscores the varying levels of redundancy and information content in the DOCI filters across different tissue classifications, offering potential pathways for optimizing imaging protocols without sacrificing model performance.

## 4. Discussion

Thyroid cancer presents a significant global health challenge due to complex anatomy, diverse etiologies, and the need for precise balance between survival outcomes and functional preservation. DOCI has emerged as a promising technology that addresses many of these challenges, offering real-time, non-invasive imaging capable of distinguishing between cancerous and non-cancerous tissues with high accuracy.

This study highlights the potential of DOCI as an effective tool in thyroid cancer management, integrating advanced imaging capabilities with cutting-edge analytical techniques. By combining DOCI with PCA, k-NN classification, and a U-Net-based segmentation model, this approach achieves remarkable accuracy in identifying and delineating tumor margins. The inclusion of feature selection and multi-filter analysis further enhances the precision of tumor characterization and segmentation, establishing a proof of concept for automating DOCI-based tumor segmentation in future applications.

The results underscore DOCI’s ability to reliably differentiate tissue types, even in cases of heterogeneous cancer presentations, providing a robust framework for both classification and segmentation tasks. The ability to operate without contrast agents, coupled with its broad field of view and real-time imaging capabilities, positions DOCI as a practical and effective intraoperative tool for tumor margin delineation. In this study, we demonstrate DOCI’s ability to recognize and differentiate thyroid cancer, which clinically can be applied in examination of FNA thyroid biopsies; in this scenario DOCI would allow for cancer detection in real time, while greatly improving the accuracy of diagnostic results.

Future work should focus on expanding DOCI’s application to larger datasets, optimizing the integration of machine learning models, and validating its clinical utility in FNA thyroid biopsies through further ex vivo studies. With continued development, DOCI has the potential to advance the diagnosis, treatment planning, and surgical outcomes for patients with head and neck cancers, ultimately improving survival rates and quality of life for patients.

## 5. Conclusion

Machine learning integrated DOCI is a potentially transformative combination of tools for the improved diagnosis and management of thyroid cancer. The ability of DOCI and machine learning to provide label-free imaging and real time analysis highlights their practical utility in clinical settings, particularly for enhancing FNA biopsy diagnostics and guiding intraoperative decision-making. This research lays the foundation for improving thyroid cancer detection, enabling precise treatment planning, and advancing patient outcomes through enhanced imaging and analysis capabilities.

## References

1. Kilfoy, B.A., et al., International patterns and trends in thyroid cancer incidence, 1973–2002. Cancer causes & control, 2009. 20: p. 525–531.

2. Pellegriti, G., et al., Worldwide increasing incidence of thyroid cancer: update on epidemiology and risk factors. Journal of cancer epidemiology, 2013. 2013(1): p. 965212.

3. Crnčić, T.B., M.I. Tomaš, N. Girotto, and S.G. Ivanković, Risk factors for thyroid cancer: what do we know so far? Acta Clinica Croatica, 2020. 59(Suppl 1): p. 66.

4. Wang, L.Y. and I. Ganly, Nodal metastases in thyroid cancer: prognostic implications and management. Future Oncology, 2016. 12(7): p. 981–994.

5. Sajisevi, M., et al., Evaluating the rising incidence of thyroid cancer and thyroid nodule detection modes: a multinational, multi-institutional analysis. JAMA Otolaryngology–Head & Neck Surgery, 2022. 148(9): p. 811–818.

6. Davies, L. and H.G. Welch, Increasing incidence of thyroid cancer in the United States, 1973-2002. Jama, 2006. 295(18): p. 2164–2167.

7. Iglesias, M.L., et al., Radiation exposure and thyroid cancer: a review. Archives of endocrinology and metabolism, 2017. 61(2): p. 180–187.

8. Brauckhoff, K. and M. Biermann, Multimodal imaging of thyroid cancer. Current Opinion in Endocrinology, Diabetes and Obesity, 2020. 27(5): p. 335–344.

9. AlSaedi, A.H., D.S. Almalki, and R.M. ElKady, Approach to Thyroid Nodules: Diagnosis and Treatment. Cureus, 2024. 16(1).

10. Lan, L., et al., Comparison of diagnostic accuracy of thyroid cancer with ultrasound-guided fine-needle aspiration and core-needle biopsy: a systematic review and meta-analysis. Frontiers in endocrinology, 2020. 11: p. 44.

11. Sanabria, A., et al., Microscopically positive surgical margins and local recurrence in thyroid cancer. A meta-analysis. European Journal of Surgical Oncology, 2019. 45(8): p. 1310–1316.

12. Kitahara, C.M. and J.A. Sosa, The changing incidence of thyroid cancer. Nature Reviews Endocrinology, 2016. 12(11): p. 646–653.

13. Tajudeen, B.A., et al., Dynamic optical contrast imaging as a novel modality for rapidly distinguishing head and neck squamous cell carcinoma from surrounding normal tissue. Cancer, 2017. 123(5): p. 879–886.

14. Kim, I.A., et al., Dynamic optical contrast imaging: A technique to differentiate parathyroid tissue from surrounding tissues. Otolaryngology–Head and Neck Surgery, 2017. 156(3): p. 480–483.

15. Pellionisz, P.A., K.W. Badran, W.S. Grundfest, and M.A.S. John, Detection of surgical margins in oral cavity cancer: the role of dynamic optical contrast imaging. Current opinion in otolaryngology & head and neck surgery, 2018. 26(2): p. 102–107.

16. Hu, Y., et al., A tool to locate parathyroid glands using dynamic optical contrast imaging. The Laryngoscope, 2021. 131(10): p. 2391–2397.

17. Tam, K., et al., Label-free, real-time detection of perineural invasion and cancer margins in a murine model of head and neck cancer surgery. Scientific Reports, 2022. 12(1): p. 12871.

18. Huang, S., et al., Ex vivo hypercellular parathyroid gland differentiation using dynamic optical contrast imaging (DOCI). Biomedical Optics Express, 2022. 13(2): p. 549–558.

19. Chatakondu, S. and K. Zhai, An Analysis of the k-Nearest Neighbor Classifier to Predict Benign and Malignant Breast Cancer Tumors. Journal of Student Research, 2023. 12(4).

20. Zhou, L., C. Wu, Y. Chen, and Z. Zhang, Multitask connected U-Net: automatic lung cancer segmentation from CT images using PET knowledge guidance. Frontiers in Artificial Intelligence, 2024. 7: p. 1423535.

21. Song, Y., P.J. Schreier, D. Ramírez, and T. Hasija, Canonical correlation analysis of high-dimensional data with very small sample support. Signal Processing, 2016. 128: p. 449–458.

22. Han, Y. and I. Joe, Enhancing Machine Learning Models Through PCA, SMOTE-ENN, and Stochastic Weighted Averaging. Applied Sciences, 2024. 14(21): p. 9772.

