## Supplementary for "Machine Learning-Based Tumor Segmentation and Classification Using Dynamic Optical Contrast Imaging (DOCI) for Thyroid Cancer"

Supplementary Figures:

| SUPPLEMENTARY TABLE 1. DOCI FILTERS | |
|---|---|
| **Filter** | **Wavelength** |
| **Filter 1** | 400nm/Long Pass |
| **Filter 2** | 413nm/10nm |
| **Filter 3** | 420nm/10nm |
| **Filter 4** | 430nm/10nm |
| **Filter 5** | 440nm/10nm |
| **Filter 6** | 450nm/10nm |
| **Filter 7** | 460nm/10nm |
| **Filter 8** | 467nm/10nm |
| **Filter 9** | 470nm/10nm |
| **Filter 10** | 473nm/10nm |
| **Filter 11** | 480nm/10nm |
| **Filter 12** | 486nm/10nm |
| **Filter 13** | 488nm/10nm |
| **Filter 14** | 492nm/10nm |
| **Filter 15** | 500nm/10nm |
| **Filter 16** | 510nm/10nm |
| **Filter 17** | 520nm/10nm |
| **Filter 18** | 532nm/10nm |
| **Filter 19** | 546nm/10nm |
| **Filter 20** | 560nm/10nm |
| **Filter 21** | 580nm/10nm |
| **Filter 22** | 589nm/10nm |
| **Filter 23** | 594nm/10nm |

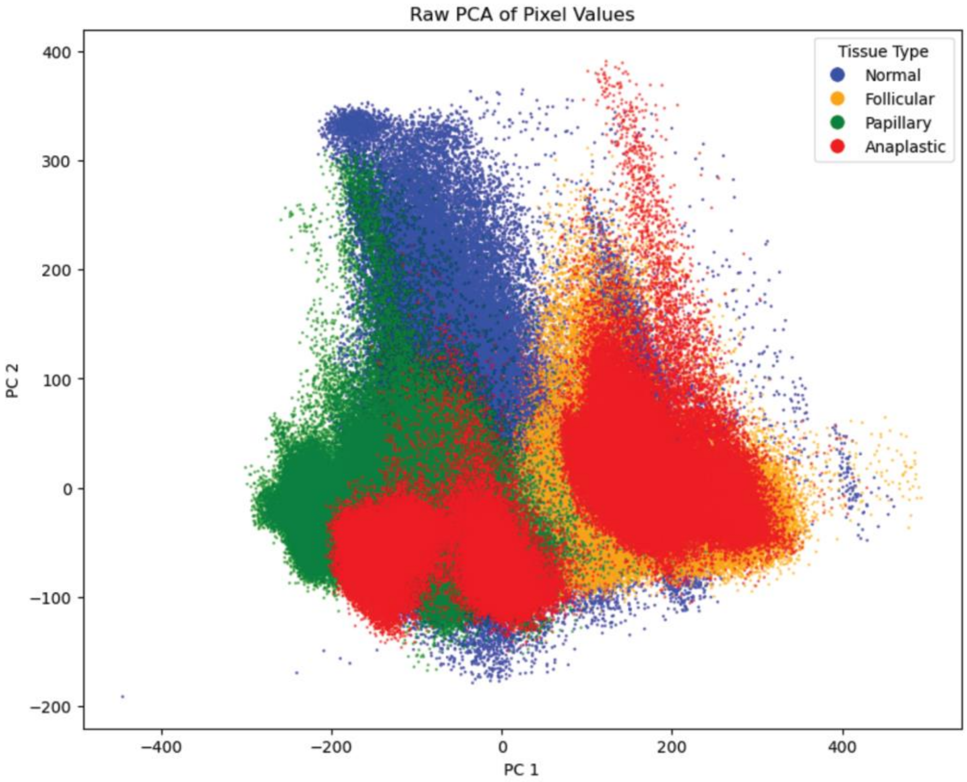


Supplementary Fig.1. PCA plot of pixel-level DOCI data, visualizing the distribution of tissue types in a two-dimensional space. Each point represents a pixel, color-coded by tissue type: Normal (blue), Follicular (yellow), Papillary (green), and Anaplastic (red). The clustering patterns highlight distinct groupings for Normal, Follicular, and Papillary tissues, while Anaplastic samples exhibit overlapping distributions, reflecting their heterogeneous nature.

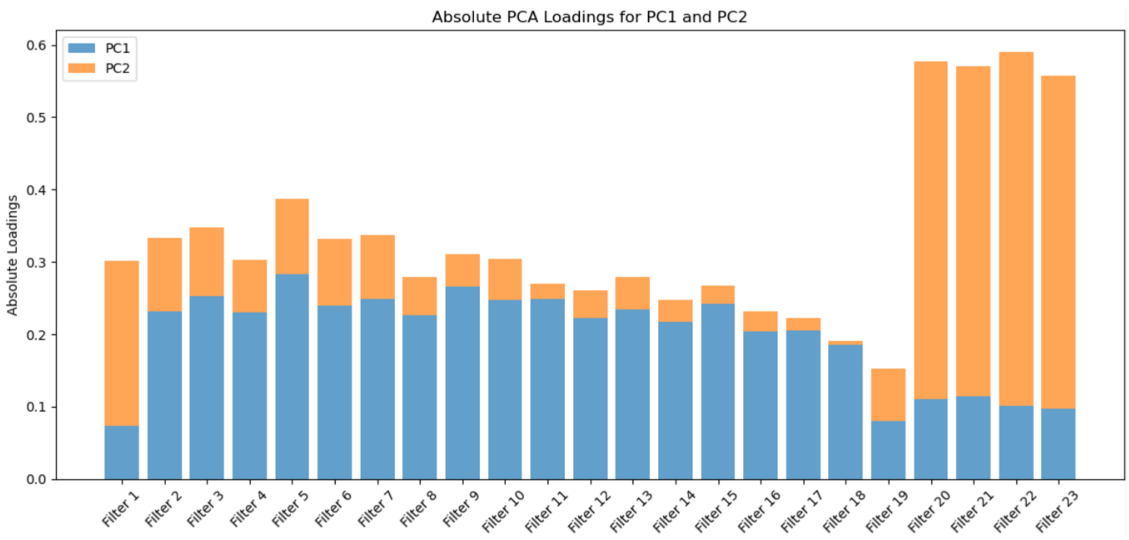


Supplementary Fig. 2. Visualization of the absolute contributions of each filter to the first two principal components (PC1 in blue and PC2 in orange).
